# A novel route for valerolactam production in *Corynebacterium glutamicum*: metabolic engineering and bioprocess optimization

**DOI:** 10.64898/2026.09.13.751229

**Authors:** Henriette Victoria Bostad, Luciana Fernandes Brito, Fernando Pérez-García

## Abstract

Valerolactam is a promising bio-based monomer for polyamide synthesis, but microbial production remains limited by inefficient 5-aminovalerate (5AVA) cyclization, by-product formation, and insufficient process optimization. In this study, *Corynebacterium glutamicum* was engineered for valerolactam production by expression of *davBA* from *Pseudomonas putida* and *lysP* from *Escherichia coli*, followed by conversion of 5AVA to valerolactam using the recently identified *avaC* gene from *Collinsella intestinalis*. Flask cultivations confirmed efficient valerolactam formation with only minor accumulation of the by-products L-lysine, 5AVA, and glutarate. Batch bioreactor experiments showed that higher glucose concentrations increased titers but also promoted by-product accumulation, whereas increasing the dissolved oxygen setpoint from 30% to 50% improved growth and volumetric productivity. Intracellular cofactor analysis revealed declining energy status and shifts in redox balance during production. Based on these findings, carbon-limited fed-batch cultivation at 50% rDO improved production performance, reaching 3.6 g/L valerolactam with a yield of 0.231 g/g and a volumetric productivity of 0.075 g/L/h, while minimizing by-product formation. These results establish AvaC-based *C. glutamicum* as a promising platform for sustainable valerolactam biosynthesis.

## 1. Introduction

The transition from fossil-based chemical manufacturing to sustainable biotechnological production routes is a central challenge in the development of the circular bioeconomy. In this context, microbial cell factories offer attractive opportunities for producing industrial platform chemicals from renewable carbon sources under mild process conditions (Jang *et al*., 2012; Yan *et al*.). Among these compounds, lactams are of particular interest because they serve as key monomers for the synthesis of polyamides, an important class of polymers used in fibers, engineering plastics, films, and other high-performance materials (Chae *et al*., 2017; Radzik *et al*., 2020). While ε-caprolactam is the established precursor of nylon-6 (Thomas and Raja, 2005), shorter-chain lactams such as the C5 δ-valerolactam (valerolactam) have gained increasing attention as precursors for nylon-5 and nylon-6,5, as well as for the development of novel bio-based polyamide materials (Han and Lee, 2023).

Valerolactam is traditionally produced through petrochemically based chemical routes that rely on fossil-derived substrates and involve harsh chemicals or reaction conditions (Gordillo Sierra and Alper, 2020). These limitations have motivated the development of alternative biosynthetic routes based on microbial production. A promising strategy involves the conversion of central metabolites into 5-aminovalerate (5AVA), followed by enzymatic activation and spontaneous or enzyme-assisted cyclization to valerolactam (Zhao *et al*., 2023). Previous studies have demonstrated the feasibility of producing 5AVA and valerolactam in engineered microbial hosts, particularly through pathways derived from L-lysine catabolism and related ω-amino-acid metabolism (Chae *et al*., 2017; Gordillo Sierra and Alper, 2020; Zhao *et al*., 2023). However, microbial valerolactam production is still limited by insufficient precursor supply, low catalytic efficiency in the cyclization of 5AVA, and competing metabolic pathways that divert carbon into unwanted by-products (Gordillo Sierra and Alper, 2020; Zhao *et al*., 2023). In addition, the roles of redox and energy states in valerolactam production remain largely unexplored and may further constrain production under certain cultivation conditions.

*Corynebacterium glutamicum* is an especially attractive platform organism for the production of nitrogen-containing chemicals (Wolf *et al*., 2021; Wendisch *et al*., 2022). This Gram-positive bacterium has a long history of safe industrial use for amino acid production and has been extensively engineered for the biosynthesis of L-lysine and other amino-acid-derived products (Wolf *et al*., 2021). Because 5AVA can be derived from L-lysine, the long-standing industrial use of *C. glutamicum* for L-lysine production, together with the availability of advanced metabolic engineering tools, makes this organism an attractive platform for the production of 5AVA and downstream valerolactam (Rohles *et al*., 2016; Jorge *et al*., 2017; Zhao *et al*., 2023). Nevertheless, further exploration of alternative biosynthetic pathways and continued bioprocess optimization will be necessary to enhance the conversion of L-lysine to valerolactam and minimize the accumulation of competing metabolites.

In this study, *C. glutamicum* was engineered for efficient 5AVA production by expression of *davBA* from *Pseudomonas putida* and *lysP* from *Escherichia coli* (Revelles *et al*., 2005; Ruiz *et al*., 2011), followed by conversion of 5AVA to valerolactam using the novel gene *avaC* from *Collinsella intestinalis*, which encodes a recently identified 5AVA cyclase with promising potential for microbial lactam biosynthesis (Zhou and Feng, 2024) (Fig. 1). To the best of our knowledge, this gene has not previously been applied in any other biotechnological production process beyond the present application. Flask cultivations showed that the major accumulated product was valerolactam, with only small amounts of the by-products 5AVA, and glutarate.

**Fig. 1:**
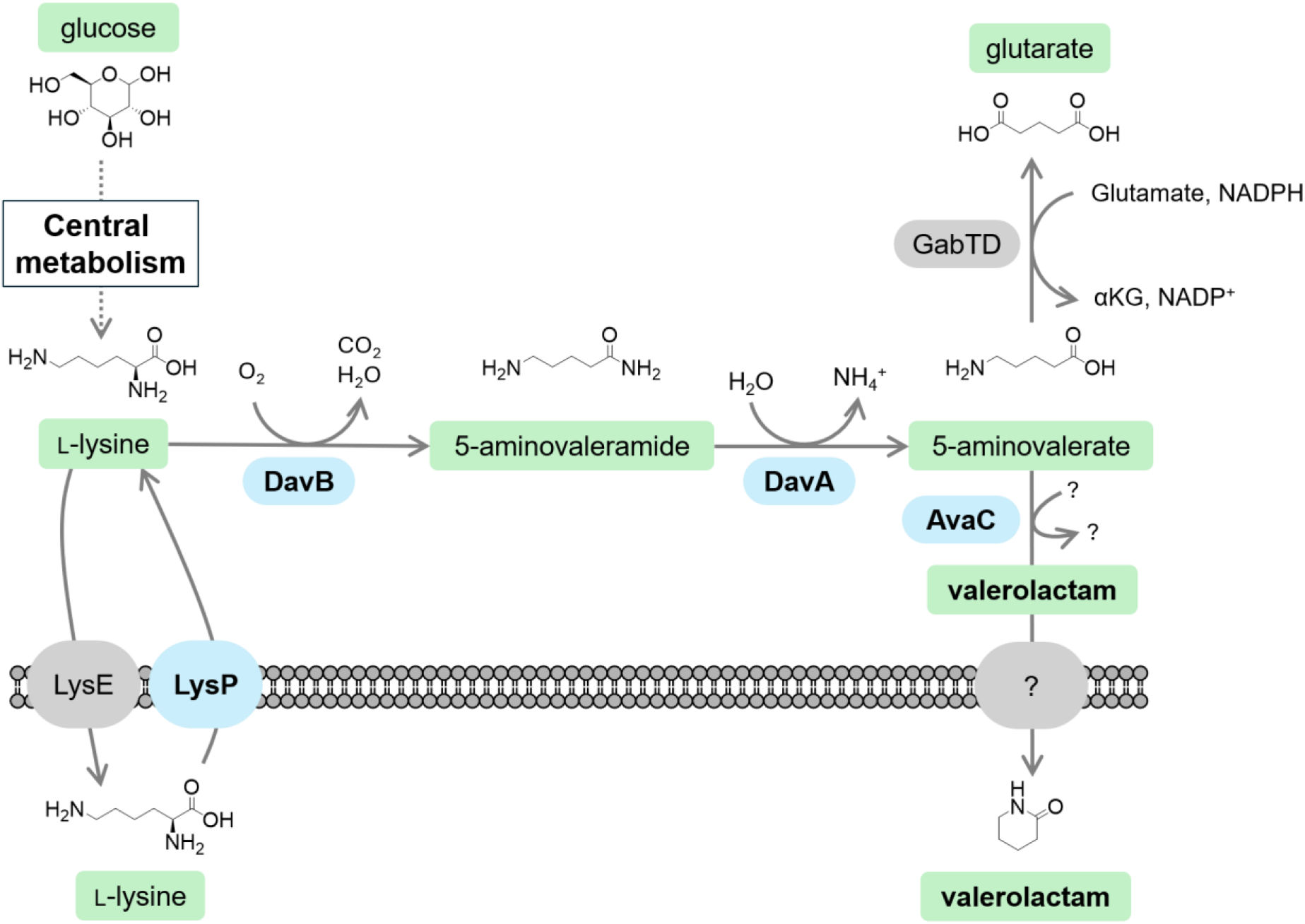
Schematic representation of the metabolic pathway used in this study for the conversion of L-lysine to 5AVA and subsequently to valerolactam, including the native competing pathway leading to the degradation of 5-aminovalerate (5AVA) to glutarate formation. Blue-shadowed elements depict non-native enzymes and transport proteins; grey-shadowed elements depict native enzymes and transport proteins. DavB, lysine 2-monooxygenase encoded by *davB* from *P. putida;* DavA, 5-aminovaleramidase encoded by *davA* from *P. putida*; AvaC, 5AVA cyclase encoded by *avaC* from *C. intestinalis*; LysP, lysine-specific permease from *E. coli*; LysE, native lysine exporter of *C. glutamicum*; GabT, native GABA/5AVA aminotransferase of *C. glutamicum*; GabD, native succinate/glutarate semialdehyde dehydrogenase of *C. glutamicum*.

Bench-scale bioreactor experiments under varying glucose and dissolved oxygen conditions highlighted the main limitations related to overflow metabolism and intracellular cofactor imbalances. This information was taken into consideration to perform optimized carbon-limited fed-batch fermentations, leading to production values, in terms of titer, yield, and volumetric productivity, that compare favorably with previously reported microbial valerolactam cell factories. These results establish a new enzymatic component for microbial lactam biosynthesis and provide a promising basis for further optimization of *C. glutamicum* as a sustainable platform for bio-based polyamide monomer production.

## 2. Materials and methods

### 2.1 Growth conditions

The plasmids and bacterial strains used in this work are listed in Table 1. Unless stated otherwise, chemicals and consumables used in this study were purchased from Sigma-Aldrich. *E. coli* DH5α was used as the cloning host and cultivated in Lysogeny Broth (10 g/L tryptone, 5 g/L yeast extract, and 5 g/L NaCl) at 37 °C and 225 rpm, either in liquid culture or on LB agar plates. *C. glutamicum* was used as the expression host and grown for pre-cultures in 2TY medium (16 g/L tryptone, 10 g/L yeast extract, and 5 g/L NaCl) at 30 °C and 225 rpm. Shake-flask cultivations were performed in baffled 250 or 500 mL flasks with a working volume corresponding to 10% of the flask volume and incubated at 30 °C and 150 rpm.

**Table 1:**
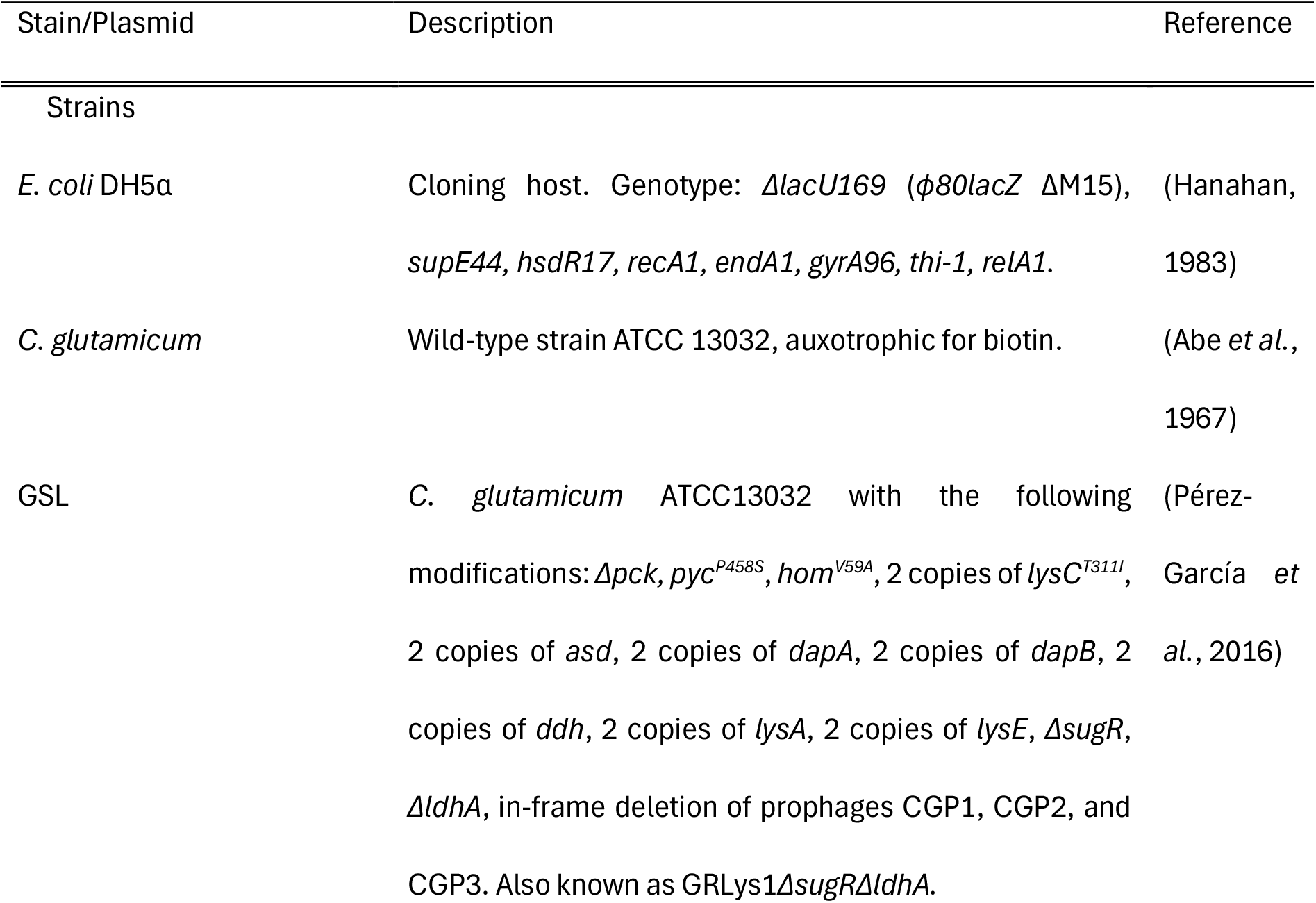
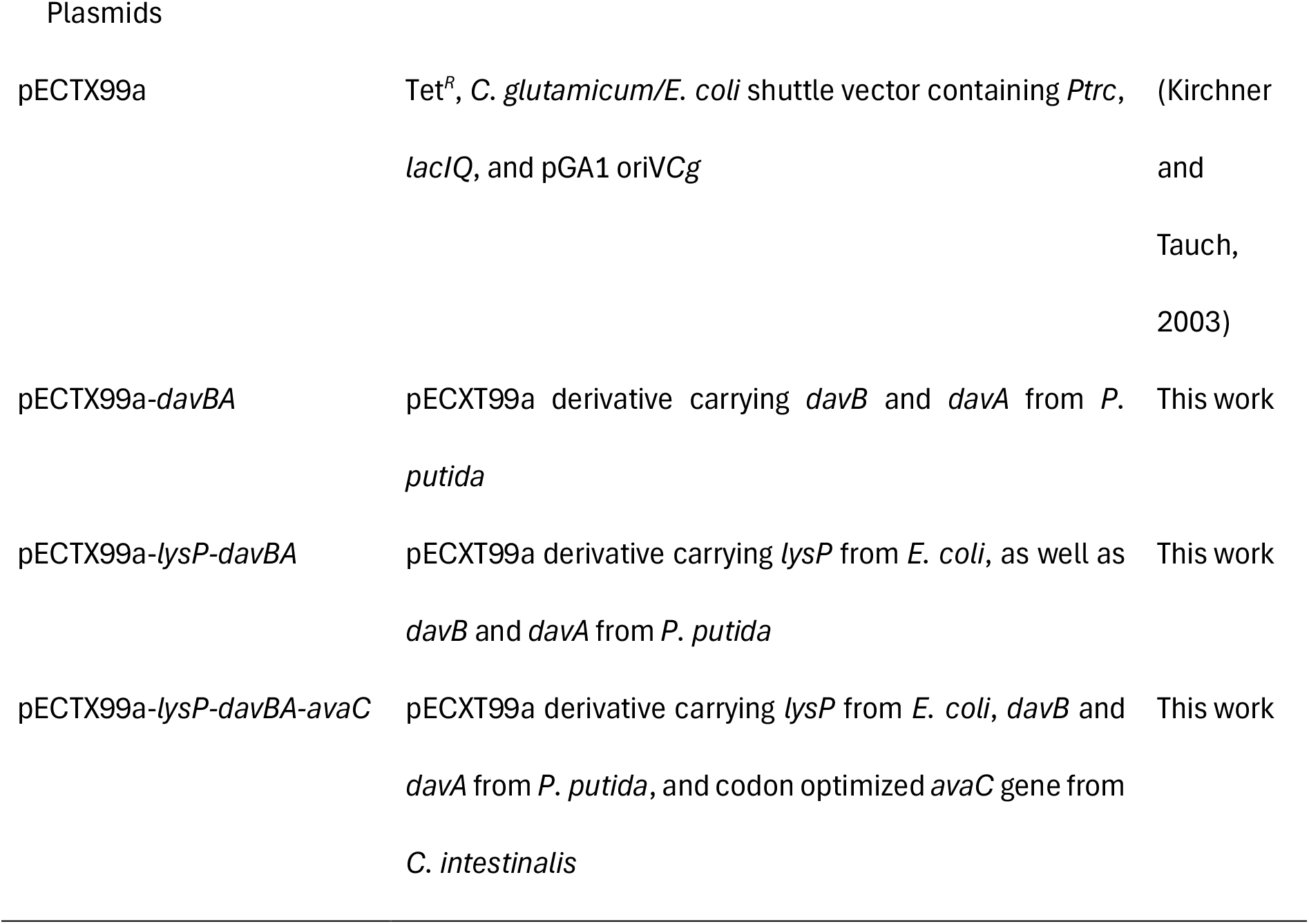
Microbial strains and plasmids used in this study.

| Stain/Plasmid | Description | Reference |
| --- | --- | --- |
| Strains |  |  |
| *E. coli* DH5α | Cloning host. Genotype: $\Delta lacU169$ ($\phi 80 lacZ$ $\Delta M15$), *supE44*, *hsdR17*, *recA1*, *endA1*, *gyrA96*, *thi-1*, *relA1*. | (Hanahan, 1983) |
| *C. glutamicum* | Wild-type strain ATCC 13032, auxotrophic for biotin. | (Abe *et al.*, 1967) |
| GSL | *C. glutamicum* ATCC13032 with the following modifications: $\Delta pck$, *pyc*<sup>P458S</sup>, *hom*<sup>V59A</sup>, 2 copies of *lysC*<sup>T311I</sup>, 2 copies of *asd*, 2 copies of *dapA*, 2 copies of *dapB*, 2 copies of *ddh*, 2 copies of *lysA*, 2 copies of *lysE*, $\Delta sugR$, $\Delta ldhA$, in-frame deletion of prophages CGP1, CGP2, and CGP3. Also known as GRLys1$\Delta sugR\Delta ldhA$. | (Pérez-García *et al.*, 2016) |
| Plasmids |  |  |
| pECTX99a | Tet<sup>R</sup>, *C. glutamicum*/*E. coli* shuttle vector containing *P<sub>trc</sub>*, *lacIQ*, and pGA1 oriVCg | (Kirchner and Tauch, 2003) |
| pECTX99a-*davBA* | pECTX99a derivative carrying *davB* and *davA* from *P. putida* | This work |
| pECTX99a-*lysP-davBA* | pECTX99a derivative carrying *lysP* from *E. coli*, as well as *davB* and *davA* from *P. putida* | This work |
| pECTX99a-*lysP-davBA-avaC* | pECTX99a derivative carrying *lysP* from *E. coli*, *davB* and *davA* from *P. putida*, and codon optimized *avaC* gene from *C. intestinalis* | This work |

Microbioreactor experiments were conducted in a BioLectorPro system (m2p Labs) using 48-well FlowerPlates sealed with gas-permeable membranes (Beckman Coulter). Each well contained 1 mL of culture, and plates were incubated at 30 °C and 1,100 rpm. The minimal medium CGXII was prepared as previously described (Eggeling *et al*., 2005) and supplemented with 0.2 g/L biotin and 1 mL/L trace metal solution. The CGXII salt solution contained 10 g/L (NH₄)₂SO₄, 1 g/L KH₂PO₄, 1 g/L K₂HPO₄, 5 g/L urea, 42 g/L MOPS, and 1 mL/L each of Mg stock solution (250 g/L MgSO₄·7H₂O) and Ca stock solution (13.25 g/L CaCl₂·2H₂O). The trace metal solution consisted of 16.4 g/L FeSO₄·7H₂O, 10 g/L MnSO₄·H₂O, 1 g/L ZnSO₄·7H₂O, 0.31 g/L CuSO₄·5H₂O, and 0.02 g/L NiCl₂·6H₂O, and was adjusted to pH 1 with HCl.

Main cultures were inoculated to an initial OD₆₀₀ of approximately 1. Optical density was determined with a Biochrom Ultrospec 7500 spectrophotometer (Fisher Scientific). Biomass concentration (g/L) was estimated according to the equation 0.343 × (OD₆₀₀ final -OD₆₀₀ initial), in which 0.343 as the lab-specific conversion factor for *C. glutamicum* (Pérez-García *et al*., 2022). Biomass yield was calculated as grams of biomass formed per gram of carbon source consumed. When indicated, media were supplemented with 1 mM IPTG, 5 µg/mL tetracycline, and/or 25 µg/mL kanamycin. All media were sterilized by autoclaving at 121 °C for 20 min. Trace metal solutions, antibiotics, biotin, and IPTG were sterilized separately by filtration through 0.22 µm pore-size filters.

### 1.2 Molecular genetics

Standard molecular biology procedures were applied as previously described (Green and Sambrook, 2012). Target genes were amplified using the CloneAmp™ HiFi PCR Premix protocol (Takara Bio Inc.). The *davA* and *davB* genes were amplified from genomic DNA of *P. putida* KT2440, whereas *lysP* was amplified from genomic DNA of *E. coli* MG1665. The *avaC* gene (*CILFYP54_00697*) from *C. intestinalis* was codon-optimized and synthesized by Integrated DNA Technologies, BV. The codon-optimized *avaC* gene sequence used in this study is provided in the Supplementary Material.

Primer sequences are listed in the Supplementary Material (Table S1). The vector pECXT99a (Kirchner and Tauch, 2003) was digested with *Bam*HI (NEB) prior to assembly. Linearized plasmids and PCR-amplified inserts were combined by Gibson assembly using the one-step isothermal DNA assembly method, with incubation at 50 °C for 1 h (Gibson *et al*., 2009). Transformation of *E. coli* was carried out by heat shock (Green and Sambrook, 2012). Colony PCR was performed with GoTaq® DNA polymerase (Promega) according to the manufacturer’s instructions. Correct plasmid construction was confirmed by Sanger sequencing (Eurofins).

Transformation of *C. glutamicum* was performed by electroporation using an Elepo21 device (Nepa Gene). Electroporation was carried out in pre-cooled cuvettes with 150 µL of competent cells and 300-1,000 ng of plasmid DNA. The following settings were used: one poring pulse at 2 kV with a pulse length of 3.5 ms, a pulse interval of 50 ms, and positive polarity, followed by three transfer pulses at 0.2 kV with a pulse length of 50 ms, a pulse interval of 50 ms, and alternating positive/negative polarity. After electroporation, cells were subjected to heat shock at 46 °C for 5 min 45 s and allowed to recover in 2TY medium for 1 h at 30 °C before plating on selective agar and incubation overnight at 30 °C.

### 1.3 Cofactors extraction

Intracellular energy related cofactors were analyzed during selected cultivation phases of the batch bioreactor experiments. Samples were taken during early exponential phase, late exponential phase and stationary phase. For each sample, 1 mL of culture was collected and immediately quenched by addition of 5 mL cold 40% methanol kept at -20◦C. The samples were centrifuged at 4,200 rpm for 6 minutes at 4◦C to remove the quenching solvent, and the resulting cell pellets were used for extraction of intracellular cofactors. Separate extraction procedures were used for adenylate metabolites, oxidized nicotinamide cofactors, and reduced nicotinamide cofactors. For extraction of ATP, ADP and AMP, the cell pellet was resuspended in 1 mL cold acetonitrile/methanol/water extraction solution (40:40:20, v/v/v) containing 0.1 M formate. The sample was vortexed thoroughly and centrifuged to clarify the extract. The supernatant was then transferred to a cold tube. For extraction of NAD^+^ and NADP^+^, the cell pellet was resuspended in 300 μL cold 0.2 M HCl, vortexed, and centrifuged at 4◦C. The supernatant was transferred to a cold tube and neutralized by addition of approximately 300 μL cold 0.1 M NaOH. The sample was vortexed thoroughly and centrifuged again to clarify the extract. For extraction of NADH and NADPH, the cell pellet was resuspended in 300 μL cold 0.2 M NaOH, vortexed, and centrifuged at 4◦C. The supernatant was transferred to a cold tube and neutralized by addition of approximately 300 μL cold 0.1 M HCl. The sample was vortexed thoroughly and centrifuged again to clarify the extract. The clarified extracts were stored at -80◦C until HPLC analysis.

### 1.4 HPLC

Extracellular compounds were quantified using a high-performance liquid chromatography (HPLC) system (Waters Alliance e2695 Separations Module). Glucose, trehalose, and lactate were analyzed on an Aminex HPX-87H column (300 × 7.8 mm; Bio-Rad) operated at 60 °C, with detection by a 2414 refractive index detector (Waters). The mobile phase consisted of 5 mM sulfuric acid applied isocratically at a flow rate of 0.6 mL/min.

For the quantification of L-lysine, L-alanine, and 5AVA, culture supernatants were derivatized with fluorenylmethoxycarbonyl chloride (FMOC) as described previously (Brito *et al*., 2021). Separation was then carried out on a Symmetry C18 column (125 × 4.6 mm, 3.5 μm; Waters) at 25 °C, and analytes were detected using a 2475 fluorescence detector (Waters). The mobile phases were buffer A (50 mM sodium acetate, pH 4.2) and buffer B (acetonitrile), applied at 1.3 mL/min with the following gradient: 0 min, 62% A / 38% B; 5 min, 62% A / 38% B; 12 min, 43% A / 57% B; 14 min, 24% A / 76% B; 15 min, 43% A / 57% B; and 18 min, 62% A / 38% B.

Valerolactam and glutarate were quantified on the same Symmetry C18 column (125 × 4.6 mm, 3.5 μm; Waters) maintained at 30 °C and detected with a 2489 UV/Vis detector (Waters) set at 210/360 nm. The mobile phase consisted of 20 mM monopotassium phosphate (pH 3) supplemented with 5% acetonitrile and was run isocratically at 1 mL/min. All extracellular HPLC analyses were performed in triplicate. ATP, ADP, AMP, NAD^+^, NADH, NADP^+^, and NADPH were quantified using a separate reversed-phase HPLC method. Samples were analyzed on a Symmetry C18 column (75 × 4.6 mm, 3.5 μm; Waters) at 25 °C. Detection was performed with a diode array detector (Agilent) at 254 nm. Mobile phase A consisted of 100 mM KH_2_PO_4_ at pH 6.0, whereas mobile phase B contained 100 mM KH_2_PO_4_, 4 mM tetrabutylammonium hydrogen sulfate (TBAHS), and 20% methanol at pH 6.0. The gradient program was as follows: 0 min A:100% B:0%; 6 min A:100% B:0%; 7.5 min A:99% B:1%; 15 min A:95% B:5%; 50 min A:20% B:80%; 55 min A:0% B:100%; 66 min A:100% B:0%.

### 1.5 Bioprocesses

Bioreactor cultivations were carried out in 2 L baffled glass reactors (Applikon Biotechnology) equipped with two 45 mm Rushton impellers positioned 6 and 12 cm above the reactor bottom. The pH and relative dissolved oxygen saturation (rDOS) were monitored using 12 mm AppliSens probes. The pH was automatically controlled at 7.0 by addition of 10% (w/w) phosphoric acid and 4 M KOH. Antifoam 204 was added manually when required. Aeration was provided at 0.75 vvm through an L-type sparger, and the cultivation temperature was maintained at 30 °C using a heating jacket.

The initial working volume during the batch phase was 0.8 L, and cultures were inoculated with overnight precultures grown in 2TY supplemented with 0.5% glucose. The batch medium consisted of modified CGXII which contained, per liter, 10 g (NH_4_)_2_SO_4_, 5 g urea, 0.5 g KH_2_PO_4_, 0.5 g K_2_HPO_4_, 0.01325 g CaCl_2_·2H_2_O, 0.25 g MgSO_4_·7H_2_O, 0.2 mg biotin, 0.005 g tetracycline, 0.083 g IPTG, and 1 mL trace element solution consisting of FeSO_4_·7H_2_O (0.025 g/L), MnSO_4_·H_2_O (0.015 g/L), ZnSO_4_·7H_2_O (0.025 g/L), CuSO_4_ (0.05 mg/L), and NiCl_2_·6H_2_O (0.005 mg/L). Glucose was supplied at either 1% or 5% as the carbon source. The rDOS was automatically maintained at either 30% or 50% by adjusting the stirrer speed, with lower and upper limits of 200 and 800 rpm, respectively. Samples were withdrawn manually using Super Safe Samplers (Infors HT).

For carbon-limited fed-batch cultivation, the feeding solution contained, per liter, 40 g glucose, 0.2 mg biotin, 0.005 g tetracycline, 0.083 g IPTG, and 1 mL of trace element solution. The feed rate was manually adjusted between 0.137 and 0.096 mL/min to prevent glucose accumulation.

## 3. Results

### 3.1 Expression of *avaC* gene enables the lactamization of 5AVA into valerolactam in *C. glutamicum*

The direct precursor of valerolactam is 5AVA. In turn, 5AVA can be derived from the proteinogenic amino acid L-lysine. Therefore, to establish valerolactam production in *C. glutamicum*, the L-lysine-producing strain GSL was used as the host. Conversion of L-lysine to 5AVA was achieved through heterologous expression of the *davBA* genes from *P. putida* (Revelles *et al*., 2005). To increase intracellular L-lysine availability and thereby enhance 5AVA formation, the *E. coli lysP* gene, encoding a lysine permease, was co-expressed with *davBA*. Finally, to generate a novel *C. glutamicum*-based valerolactam producer, the *avaC* gene from *C. intestinalis* (Zhou and Feng, 2024) was co-expressed, enabling the lactamization of 5AVA to valerolactam (Figure 2). Therefore, the newly constructed strains GSL(pECTX99a-*davBA*), GSL(pECTX99a-*lysP*-*davBA*), and GSL(pECTX99a-*lysP*-*davBA-avaC*) were tested in CGXII minimal medium with 1% glucose as sole carbon source. The strain GSL(pECTX99a) were used as control strain. Growth data (Fig. 2A) and product data from supernatants collected after glucose depletion (Fig. 2B) were obtained for each strain. Under the conditions tested, no significant differences among the tested strains were observed with regard growth rates and biomass yield values (Fig. 2A). As expected, the major product observed in the supernatants of GSL(pECTX99a) was L-lysine reaching a titer of 1.9 ± 0.4 g/L. on the other hand, the strain GSL(pECTX99a-*davBA*) produced a mixture of L-lysine (0.8 ± 0.3 g/L), 5AVA (0.6 ± 0.1 g/L), and its degradation product glutarate (0.1 ± 0.0 g/L). The strain GSL(pECTX99a-*lysP*-*davBA*) showed an increase in 5AVA (1.4 ± 0.3 g/L), and glutarate (0.3 ± 0.1 g/L) titers, but no L-lysine was observed in the supernatants. Finally, the strain GSL(pECTX99a-*lysP*-*davBA-avaC*) produced 1.4 ± 0.1 g/L of valerolactam together with 0.2 ± 0.1 g/L of 5AVA and 0.1 ± 0.0 g/L of glutarate (Fig. 2B).

**Fig. 2:**
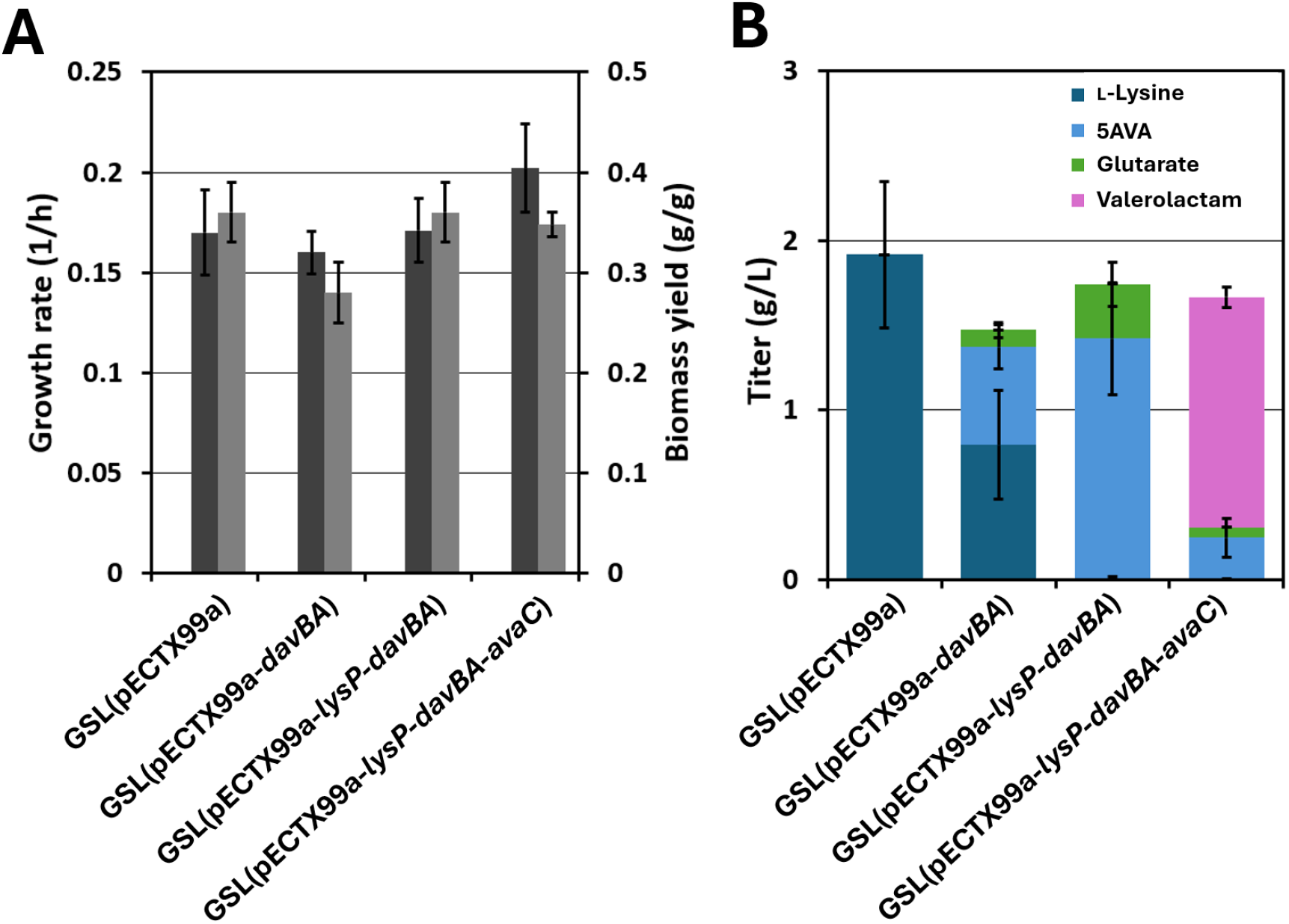
Growth parameters and titer values of the *C. glutamicum* strains used in this study, cultivated in shake flasks. **(A)** Growth rates (dark grey bars) and biomass yield values (light grey bars). **(B)** L-Lysine, 5AVA, glutarate, and valerolactam titers. Average values and standard deviations of triplicates are shown.

These results proved that *avaC* gene from *C. intestinalis* can efficiently synthesize valerolactam from 5AVA in *C. glutamicum*-based cell-factories.

### 3.2 Effects of 5AVA and valerolactam accumulation on *C. glutamicum* growth and biomass formation

Accumulation of pathway products and intermediates can limit microbial production by affecting cell growth. For L-lysine-derived valerolactam production, this is relevant because engineered *C. glutamicum* cells are exposed to both the intermediate 5AVA and the final product valerolactam. Therefore, wild-type *C. glutamicum* was cultivated in a BioLector in 2TY complex medium supplemented with 1% glucose and increasing concentrations of valerolactam, ranging from 0 to 200 mM. Growth rates and final biomass values were determined (Fig. 3A), showing no significant differences in final biomass values but a decrease in growth rates with increasing valerolactam concentrations, resulting in an estimated inhibitory constant (*Ki*) of 165 mM (Fig. 3A). Similarly, wild-type *C. glutamicum* was also cultivated in a BioLector in 2TY complex medium supplemented with 1% glucose, 200 mM 5AVA, and increasing concentrations of valerolactam (0 to 200 mM) (Fig. 3B). Under these conditions the estimated *Ki* value was 208 mM of valerolactam. Additionally, when combining 200 mM of both 5AVA and 200 mM of valerolactam there was a significant decrease in biomass formation (Fig. 3B).

**Fig. 3:**
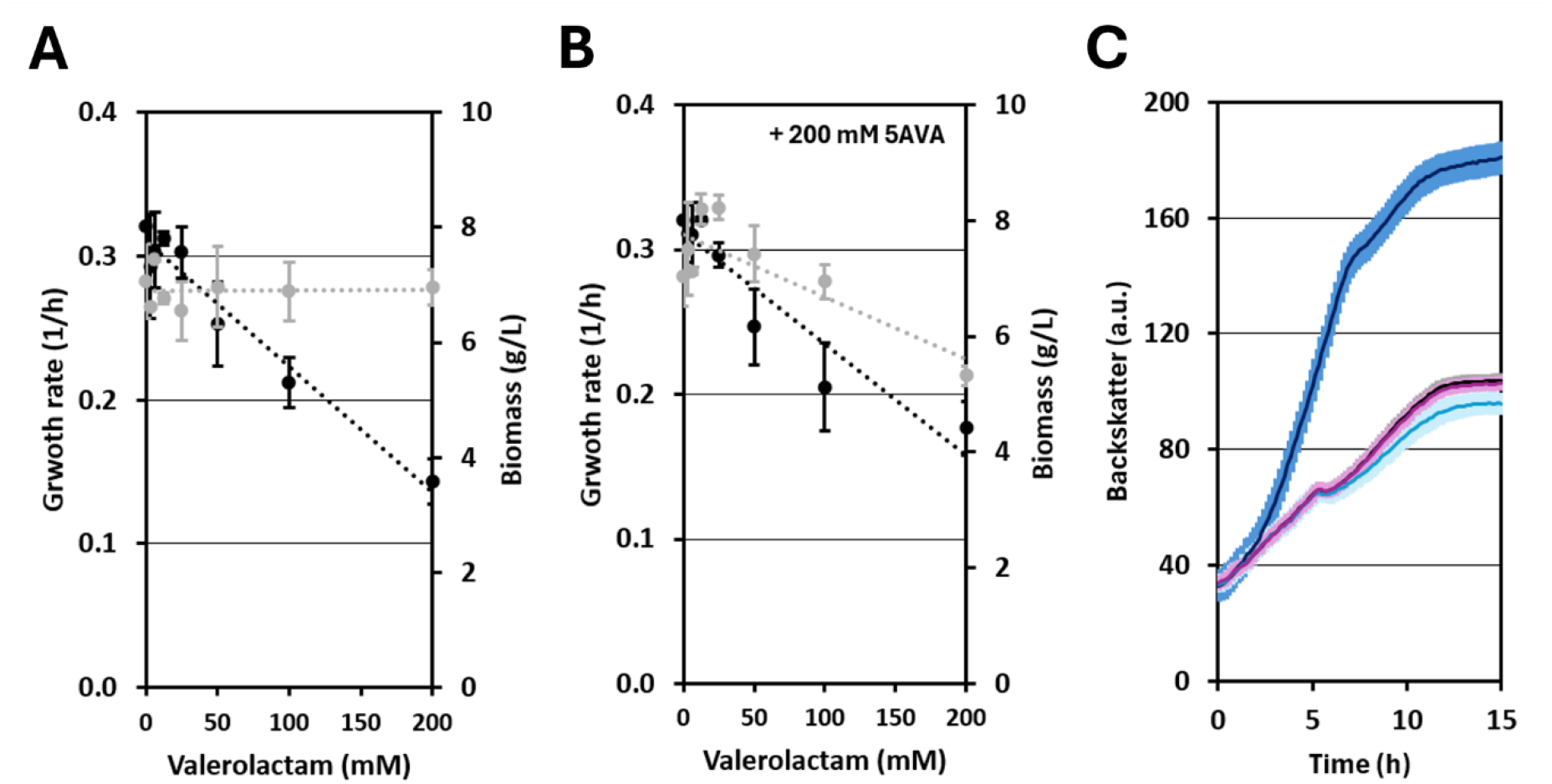
**(A)** Growth rates (black dots) and final biomass (grey dots) of *C. glutamicum* grown in 2TY complex medium supplemented with 1% glucose and increasing concentrations of valerolactam. **(B)** Growth rates (black dots) and final biomass (grey dots) of *C. glutamicum* grown in 2TY complex medium supplemented with 1% glucose, 200 mM 5AVA, and increasing concentrations of valerolactam. Trendlines are represented with dotted lines **(C)** *C. glutamicum* growth in 2TY (black) or 2TY supplemented with either 25mM glucose (dark blue), 25mM 5AVA (light blue), or 25mM Valerolactam (purple). Average values and standard deviations of biological triplicates are shown.

Furthermore, 5AVA and valerolactam were evaluated as carbon sources for *C. glutamicum*. Hence, *C. glutamicum* wild-type was cultivated in a BioLector in 2TY plus 25mM glucose, 2TY plus 25mM 5AVA, 2TY plus 25mM valerolactam, or plain 2TY as control (Fig. 3C). Under these conditions, only *C. glutamicum* growing on 2TY plus 25mM glucose showed increase in biomass formation while *C. glutamicum* growing on 2TY plus 25mM valerolactam showed no changes as compared with *C. glutamicum* growing in plain 2TY. However, *C. glutamicum* growing on 2TY plus 25mM 5AVA showed slight decrease in biomass as compared to the control.

Overall, these results indicate that *C. glutamicum* tolerates relatively high concentrations of valerolactam, although increasing valerolactam levels progressively reduce growth rate. However, 5AVA appears to impose a stronger physiological burden, particularly when combined with valerolactam, leading to reduced biomass formation and suggesting that accumulation of pathway intermediates may be more limiting than valerolactam itself.

### 3.3 Bench-bioprocesses

Bioreactor cultivations were performed to evaluate valerolactam production by the strain GSL(pECXT99a-*lysP-davBA-avaC*) under controlled conditions (Fig. 4). Four batch cultivations were carried out, combining two initial glucose concentrations, 1% and 5%, with two relative dissolved oxygen (rDO) setpoints, 30% and 50%. Biomass formation, glucose consumption, valerolactam production, dissolved oxygen, CO₂, and extracellular by-products were monitored over time. The monitored by-products included pathway intermediates such as L-lysine and 5AVA, competing metabolites such as glutarate, overflow indicators such as lactate and L-alanine, and the stress indicator trehalose (Fig. 5). Cultivations were terminated when the rDO increased above 30% and CO_2_ reached basal levels, indicating carbon source depletion.

**Fig. 4:**
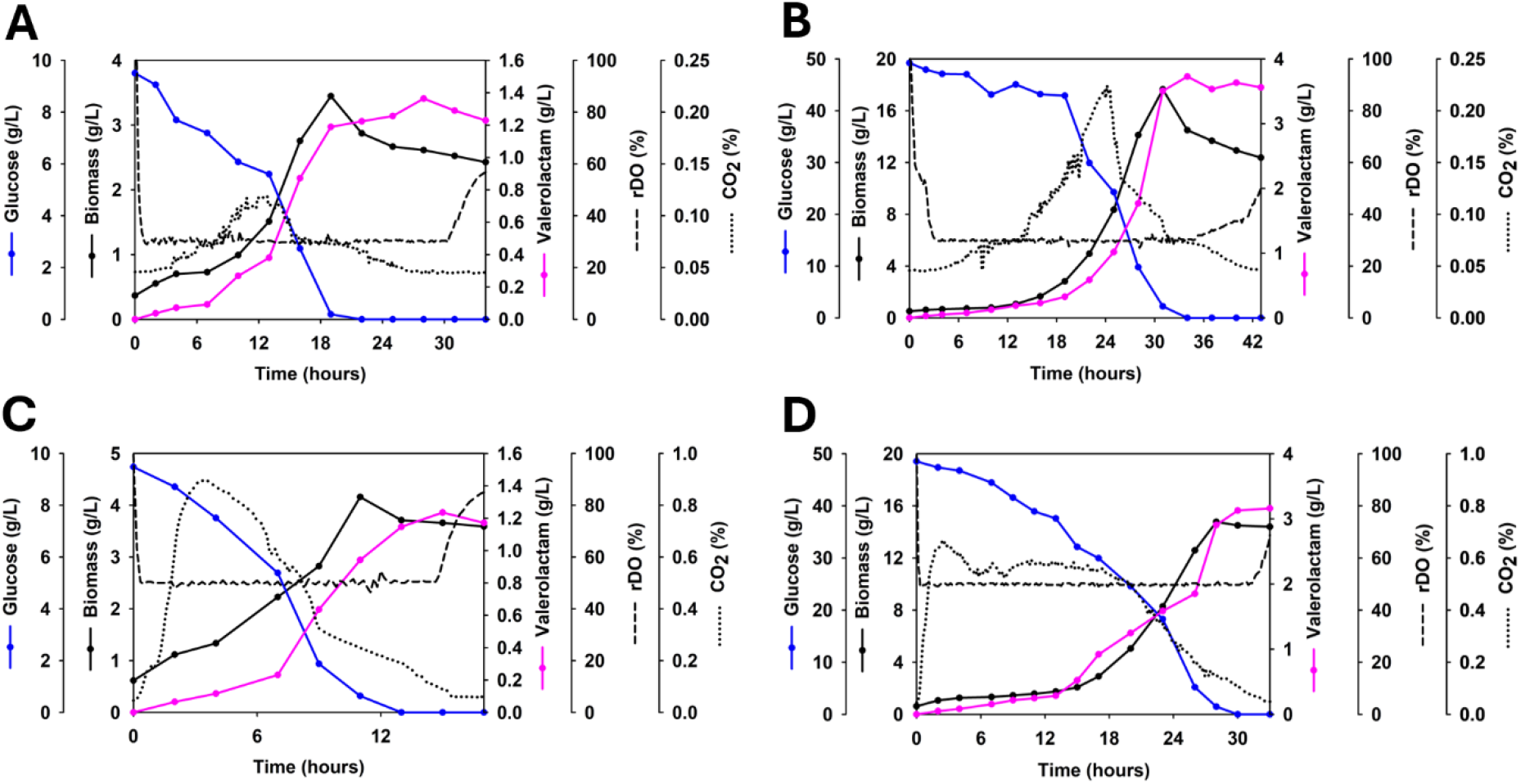
Bench-scale batch reactor cultivations of the *C. glutamicum* strain GSL(pECXT99a-*lysP-davBA-avaC*). The data shown include biomass (g/L), glucose (g/L), valerolactam (g/L), rDO (%), and off-gas CO₂ (%). **(A)** 1% glucose and 30% rDO. **(B)** 5% glucose and 30% rDO. **(C)** 1% glucose and 50% rDO. **(D)** 5% glucose and 50% rDO.

**Fig. 5:**
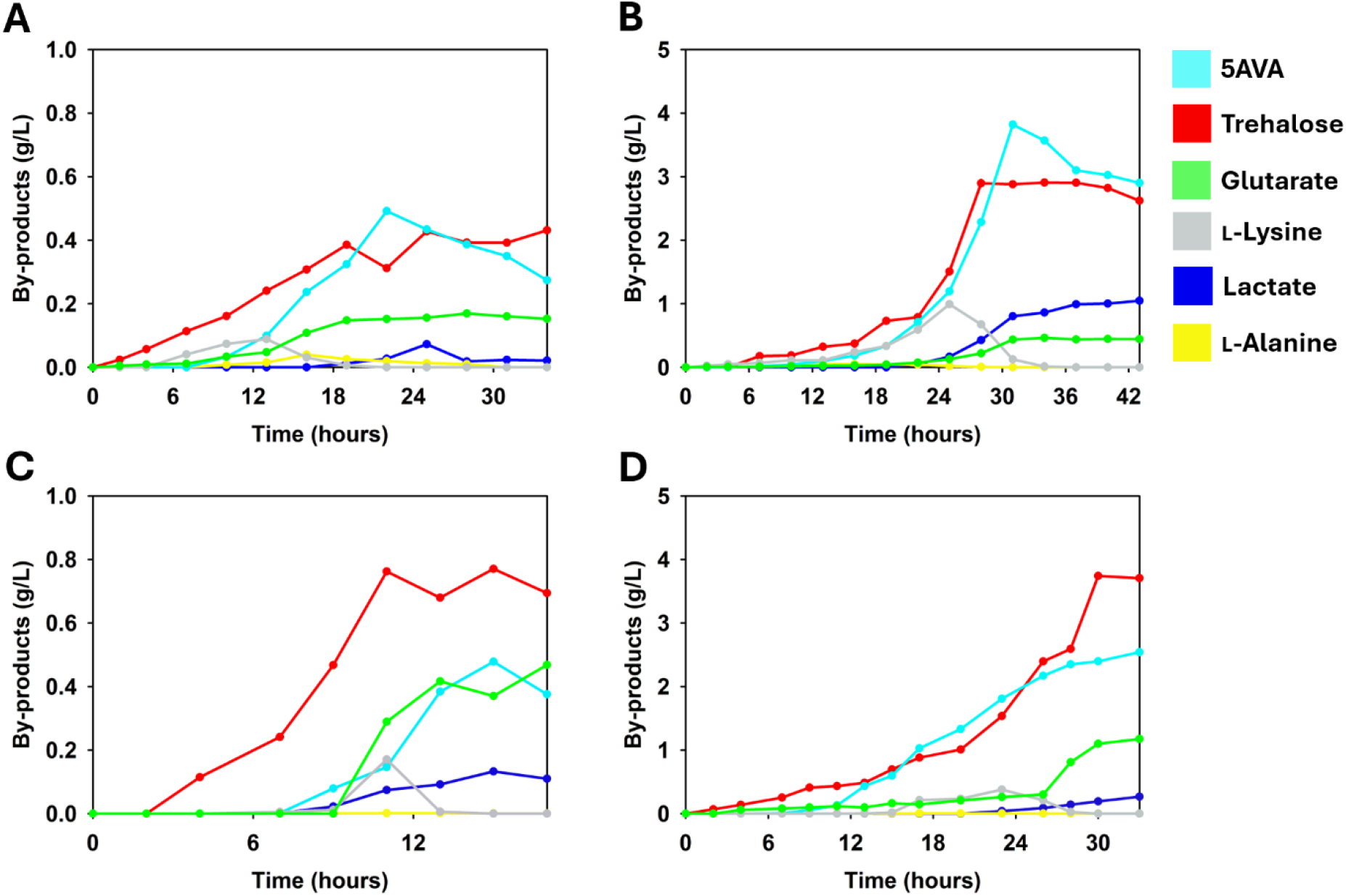
By-products formed during the bench-scale batch reactor cultivations of the *C. glutamicum* strain GSL(pECXT99a-*lysP-davBA-avaC*). **(A)** 1% glucose and 30% rDO. **(B)** 5% glucose and 30% rDO. **(C)** 1% glucose and 50% rDO. **(D)** 5% glucose and 50% rDO.

Valerolactam accumulated throughout all bioreactor cultivations. At 30% rDO, the cultivation with 1% glucose reached 1.2 g/L valerolactam after 34 h, corresponding to a yield of 0.120 g/g (Fig. 4A), whereas increasing the initial glucose concentration to 5% enhanced production, resulting in a maximum concentration of 3.6 g/L valerolactam after 43 h, corresponding to a yield of 0.072 g/g (Fig. 4B). At 50% rDO, the cultivation with 1% glucose reached 1.2 g/L valerolactam after 17 h, corresponding to a yield of 0.120 g/g (Fig. 4C), while the cultivation with 5% glucose reached 3.2 g/L valerolactam after 33 h, corresponding to a yield of 0.064 g/g (Fig. 4D).

By-product accumulation differed markedly depending on the initial glucose concentration and rDO setpoint (Fig. 5). In the 1% glucose cultivation at 30% rDO, only limited by-product accumulation was observed, with maximum concentrations of 0.3 g/L 5AVA, 0.4 g/L trehalose, and < 0.1 g/L lactate while glutarate was not detected (Fig. 5A). In contrast, the 5% glucose cultivation at 30% rDO showed higher accumulation of pathway-related by-products, including 2.9 g/L 5AVA and 0.4 g/L glutarate. Trehalose also accumulated to 2.6 g/L, and lactate increased during the later phase of cultivation, reaching 1.0 g/L (Fig. 5B). At 50% rDO, the 1% glucose cultivation showed final concentrations of 0.4 g/L 5AVA and 0.7 g/L trehalose, while glutarate increased to 0.5 g/L toward the end of cultivation. Lactate remained low, reaching a maximum concentration of 0.1 g/L (Fig. 5C). In contrast, the 5% glucose cultivation at 50% rDO again showed higher by-product accumulation, with 5AVA, trehalose, and glutarate reaching 2.5, 3.7, and 1.2 g/L, respectively. Lactate reached 0.3 g/L (Fig. 5D). L-Alanine and L-lysine were detected under all conditions and were consumed before the end of the cultivations.

Taken together, GSL(pECXT99a-*lysP-davBA-avaC*) produced valerolactam under controlled bioreactor conditions, with higher glucose concentrations increasing final titers. However, higher glucose availability also increased by-product and stress-metabolite accumulation, highlighting the need for further pathway and bioprocess optimization. Increasing the rDO setpoint from 30% to 50% improved volumetric productivity under both glucose concentrations, mainly by shortening the cultivation time.

### 3.4 Valerolactam production is associated with changes in cellular energy and redox states

To evaluate the cellular energy and redox states during valerolactam production, intracellular cofactors were analyzed. Samples for cofactor extraction were collected from batch bioreactor cultivations during the exponential, late exponential, and stationary phases of cells grown with either 1% or 5% glucose at rDO setpoints of 30% or 50%. Cofactor extractions were performed as technical duplicates and included ATP, ADP, AMP, NADH, NADPH, NAD⁺, and NADP⁺. These data were used to calculate the ATP/ADP ratio, adenylate energy charge (AEC), NAD⁺/NADH ratio, and NADP⁺/NADPH ratio (Fig. 6).

**Fig. 6:**
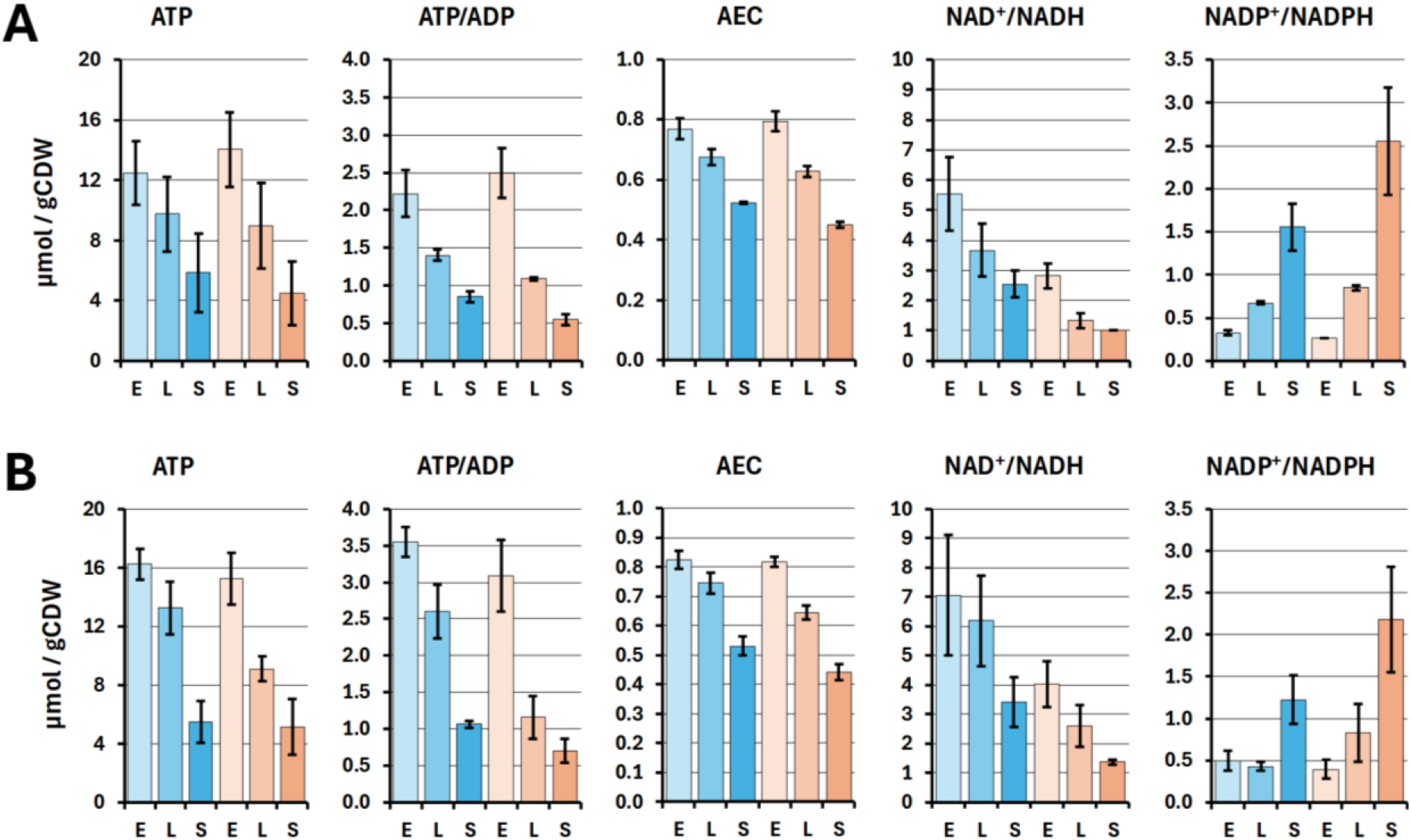
Intracellular cofactor profiles of GSL(pECXT99a-*lysP-davBA-avaC*) during batch bioreactor cultivations at 30% rDO **(A)** and 50% rDO **(B)** using 1% (Blue bars) or 5% glucose (orange bars) as carbon source. The analyzed parameters include ATP levels, ATP/ADP ratio, AEC, NAD+/NADH ratio, and NADP+/NADPH ratio in the units µmol/gCDW. Samples were collected during the early exponential phase (E), late exponential phase (L), and stationary phase (S). Average values and standard deviations of technical duplicates are shown.

Intracellular cofactor analysis showed a progressive decrease in cellular energy status during valerolactam production at both rDO setpoints. At 30% rDO, ATP levels, ATP/ADP ratios, and adenylate energy charge (AEC) decreased from the early exponential phase toward the stationary phase under both tested glucose concentrations (Fig. 6A). A similar trend was observed at 50% rDO, although with 1% glucose the early-phase ATP levels and ATP/ADP ratios were generally higher than those observed at 30% rDO also with 1% glucose (Fig. 6B). This suggests that higher oxygen availability improved the initial cellular energy state, which may contribute to the increased volumetric productivity observed at 50% rDO (Fig. 4C). The NAD⁺/NADH ratio decreased over time at both 30% and 50% rDO, indicating a shift toward a more reduced intracellular state during cultivation. This decrease was particularly evident in cultures grown with 5% glucose, suggesting that higher carbon availability increased NADH formation and contributed to redox imbalance. In contrast, the NADP⁺/NADPH ratio increased toward the stationary phase, especially under 5% glucose conditions at both rDO setpoints, indicating relative NADPH depletion or increased demand for NADPH-dependent reactions during later stages of cultivation.

Taken together, these results show that valerolactam production was associated with a progressive decrease in cellular energy status and a shift in redox balance at both 30% and 50% rDO. Higher oxygen availability improved the early cellular energy state and volumetric productivity, whereas higher glucose availability intensified redox imbalance and NADPH demand during later cultivation phases.

### 3.5 Carbon limited fed-batch

Based on the information gathered so far, a new bioreactor cultivation with the strain GSL(pECXT99a-*lysP-davBA-avaC*) was performed in carbon-limited fed-batch mode at 50% rDO (Fig. 7). The batch phase started with 1% glucose as the carbon source, and feeding was initiated after 11 hours, when the cells had reached the late-exponential phase. At that point, the glucose concentration in the fermenter vessel was 0.85 g/L. Glucose concentration was monitored by HPLC, and the feeding rate was manually adjusted between 0.137 and 0.096 mL/min to prevent glucose accumulation. By the end of the cultivation, the total feed volume was 206 mL and the total amount of glucose supplied was 15.6 g. Under these conditions, the final valerolactam titer, yield, and volumetric productivity reached 3.6 g/L, 0.231 g/g, and 0.075 g/L/h, respectively (Fig. 7A). Interestingly, the valerolactam yields and volumetric productivities differed between the cultivation phases. The valerolactam yield was 0.103 g/g during the batch phase and 0.344 g/g during the feeding phase, whereas the productivity values were 0.103 g/L/h and 0.066 g/L/h, respectively. On the other hand, only limited by-product accumulation was observed, with maximum concentrations of 0.3 g/L 5AVA, 0.5 g/L trehalose and 0.2 g/L glutarate, while L-lysine, and L-alanine were not detected under these conditions, and lactate was consumed before the end of the cultivation (Fig. 7B).

**Fig. 7:**
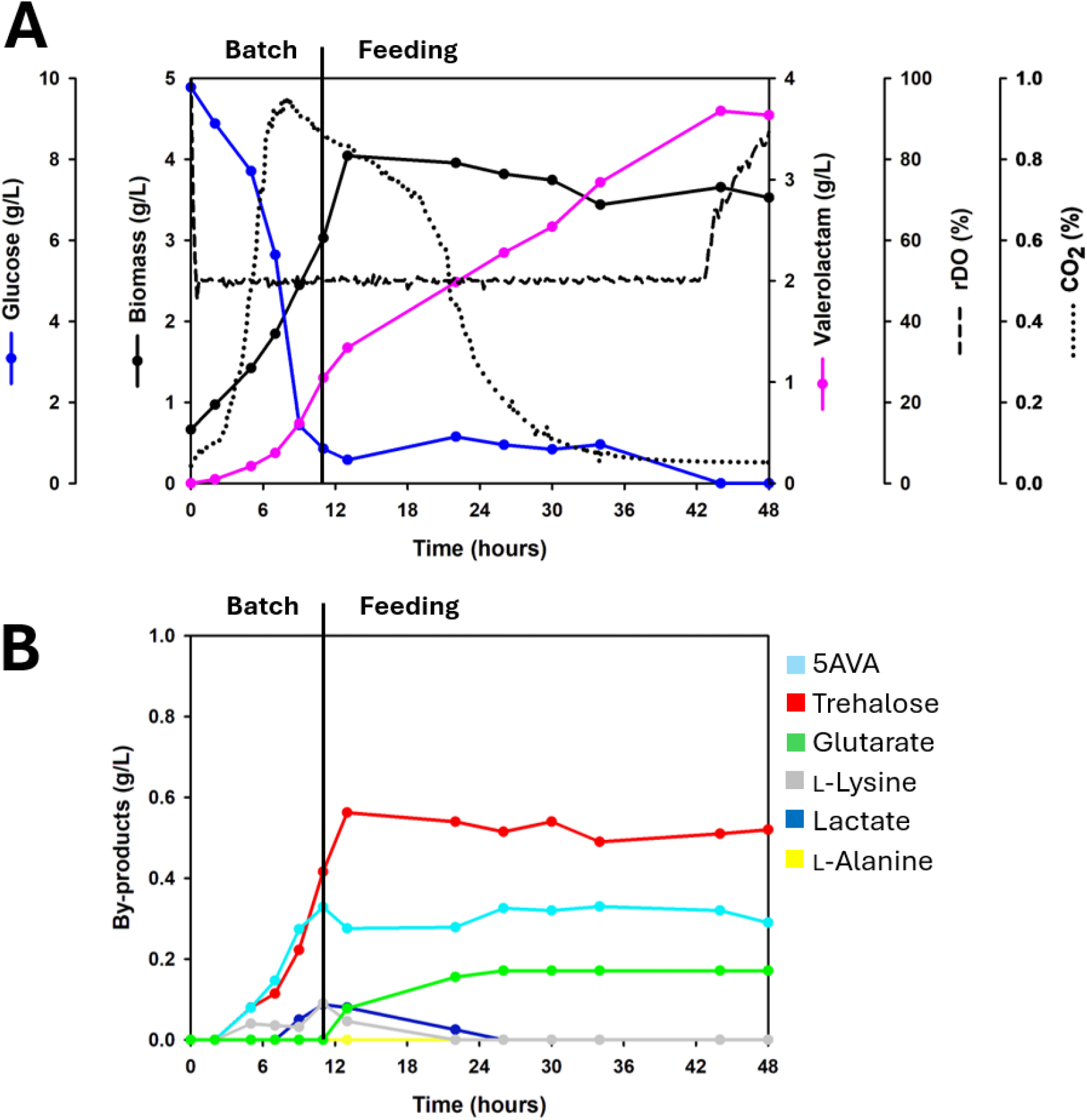
Bench-scale fed-batch reactor cultivation data of strain GSL(pECXT99a-*lysP-davBA-avaC*). **(A)** The data shown here include biomass (g/L), glucose (g/L), valerolactam (g/L), rDO (%), stirring frequency (rpm), and off-gas CO₂ (%). **(B)** By-products formed during the bench-scale fed-batch reactor cultivation.

In conclusion, glucose-based carbon-limited fed-batch cultivation at 50% rDO improved the valerolactam titer and yield of GSL(pECXT99a-*lysP-davBA-avaC*) while minimizing carbon loss to unwanted metabolites.

## 4. Discussion

*C. glutamicum* is a highly versatile bacterial cell factory that has been widely engineered for the production of amino acids, non-proteinogenic amino acids, diamines, antimicrobials, carotenoids, organic acids, and vitamins (Wolf *et al*., 2021; Christmann *et al*., 2023; Brito *et al*., 2025). More recently, lactams have been added to its product portfolio (Zhang *et al*., 2022; Han and Lee, 2023; Zhao *et al*., 2023). This study demonstrates that *avaC* gene from the human gut bacterium *C. intestinalis* (Zhou and Feng, 2024) can be used in *C. glutamicum*-based cell-factories for the lactamization of 5AVA to valerolactam. Previously, valerolactam production from 5AVA has been established for instance by expression of *act* gene encoding a β-alanine CoA transferase from *Clostridium propionicum* in *E. coli* and *C. glutamicum* (Chae *et al*., 2017; Han and Lee, 2023). The proposed reaction mechanism for lactam formation by Act begins with activation of the carboxyl group of the ω-amino acid substrate, in this case 5AVA, which is then followed by spontaneous intramolecular cyclization to form the lactam ring (Chae *et al*., 2017). Han and Lee (2023) reported progressive improvements in valerolactam production in *C. glutamicum* based of the Act enzyme and through stepwise strain engineering. First, the VL1 strain overexpressing the codon-optimized *act* gene produced 2.0 g/L valerolactam in flask cultivations with 8% glucose as the carbon source, corresponding to a yield of 0.025 g/g. Further improvement of L-lysine supply together with elimination of glutarate formation led to strain VL3, which increased valerolactam production to 3.84 g/L and the yield to 0.048 g/g. Additionally, start codon exchange from ATG to GTG of the genes *hom* and *gdh* as well as promoter exchange to enhance expression of *NCgl0464*, encoding a transport protein, which promoted 5AVA reuptake, further increased valerolactam production to 5.1 g/L, corresponding to a yield of 0.063 g/g, in flask cultivations with 8% glucose (Han and Lee, 2023). Additionally, Zhao et al. (2023) reported valerolactam production in *C. glutamicum* through codon-optimized heterologous expression of *orf26* from *Streptomyces aizunensis* and *caiC* from *E. coli* in the 5AVA-producing strain Val-1, resulting in the valerolactam-producing strains Val-10 and Val-12. In flask cultivations with 10% glucose as the carbon source, Val-10 and Val-12 produced 2.2 and 1.4 g/L valerolactam, respectively, corresponding to yields of 0.022 and 0.014 g/g (Zhao *et al*., 2023). In comparison, the strain GSL(pECTX99a-*lysP-davBA-avaC*) developed in the present study reached 1.4 g/L valerolactam from 1% glucose in flask cultivations, corresponding to a yield of 0.140 g/g (Fig. 2). While Act catalyzes the activation of 5AVA using Acyl-CoA as a coenzyme, ORF26 mediates the activation of 5AVA via ATP and CaiC utilizes CoA and ATP as cofactors (Kugler *et al*., 2020; Zhou and Feng, 2024). The protein encoded by *avaC* was identified as a 5AVA cyclase that catalyzes the dehydration and cyclization of 5AVA to valerolactam (Zhou and Feng, 2024). However, its detailed catalytic mechanism and potential cofactor or coenzyme requirements have not yet been clearly established. Therefore, although substrate-activation chemistry may be involved, it remains unclear whether AvaC directly depends on ATP, acetyl-CoA, or another activating molecule. When cultivated at 50% rDO instead of the classical 30% setpoint, GSL(pECTX99a-*lysP-davBA-avaC*) showed faster growth, which in turn resulted in higher volumetric productivity (Fig. 4). Oxygen availability may influence this process at different levels. On the one hand, the first pathway enzyme, lysine 2-monooxygenase encoded by *davB*, is oxygen dependent. It requires molecular oxygen to oxidatively decarboxylate L-lysine into 5-aminopentanamide (Revelles *et al*., 2005). On the other hand, higher oxygen availability may enhance activity of the electron transport chain, sustaining a stronger proton motive force. This can result in a higher cellular energy state, reflected by a larger ATP pool and a higher ATP/ADP ratio, thereby supporting active biomass formation and faster growth rates (Michel *et al*., 2015). Additionally, as mentioned above, the mechanism of AvaC remains uncharacterized, and it is therefore possible that it uses ATP as an activating molecule for 5AVA, similar to ORF26 from *S. aizunensis* (Zhao *et al*., 2023). High-cell-density cultivations may create microanaerobic zones when the oxygen supply is insufficient. However, this effect is more relevant in industrial bioreactors than at the scale of this study (Nadal-Rey *et al*., 2021). However, when carbon is supplied in excess and/or oxygen availability becomes limiting, *C. glutamicum* may shift toward overflow metabolism, resulting in the formation of by-products such as lactate or alanine (Inui *et al*., 2004; Wieschalka *et al*., 2012; Hasegawa *et al*., 2017). The results of this study show that, under 5% glucose conditions, lactate formation decreased at 50% rDO compared with 30% rDO (Fig. 5), suggesting that oxygen limitation contributed to lactate accumulation. Interestingly, trehalose levels also varied (Fig. 5). Trehalose can be considered a stress-associated metabolite in *C. glutamicum*, particularly in relation to osmotic stress and cell-envelope physiology (Wolf *et al*., 2003; Burkovski, 2013). Trehalose formation by *C. glutamicum* strains in reactor bioprocesses has been reported previously when overproducing 5AVA but not when overproducing the 5AVA-derivative glutarate (Rohles *et al*., 2016, 2018). The results of this study show that 5AVA and trehalose are the main by-products, and that their concentrations change in parallel under different cultivation conditions (Fig. 5). This could indicate 5AVA-derived stress. 5AVA extracellular toxicity in *C. glutamicum* has been evaluated previously reporting a *Ki* of 1.1 M under the conditions tested (Jorge *et al*., 2017). In this study, the extracellular toxicity of valerolactam, alone or in combination with 5AVA, was evaluated (Fig. 3), and no significant growth impairment was observed at the production levels achieved here. However, when 5AVA was tested as a carbon source, a decrease in final biomass was observed (Fig. 3C). This observation, together with trehalose co-production, may suggest that 5AVA imposes physiological stress on *C. glutamicum.* Taken together, these results motivated the implementation of a carbon-limited fed-batch cultivation at 50% rDO, which resulted in a valerolactam titer, yield, and volumetric productivity of 3.6 g/L, 0.231 g/g, and 0.075 g/L/h, respectively, together with 0.2 g/L 5AVA and 0.6 g/L trehalose (Fig. 7). Previous glucose-based fed-batch approaches have reported valerolactam titers of 1.2 g/L with a *E. coli* strain, as well as 12.3 and 76.1 g/L with *C. glutamicum* strains (Chae *et al*., 2017; Han and Lee, 2023; Zhao *et al*., 2023). In particular, Han and Lee, using strain VL10(pVL1), achieved the best production values reported to date, with a valerolactam titer, yield, and volumetric productivity of 76.1 g/L, 0.280 g/g, and 0.990 g/L/h, respectively (Han and Lee, 2023). Their strategy included multiple copies of the *act* gene, elimination of glutarate formation, and transport engineering to promote 5AVA reuptake. *C. glutamicum* possesses the *gabTD* genes, encoding a GABA/5AVA aminotransferase and a succinate/glutarate semialdehyde dehydrogenase, respectively (Rohles *et al*., 2016). While overexpression of the native *gabTD* in a 5AVA-producing *C. glutamicum* strain increases glutarate production (Pérez-García *et al*., 2018), disruption of these genes is commonly applied to reduce diversion of 5AVA toward glutarate (Rohles *et al*., 2016; Han and Lee, 2023). Additionally, the gene *lysE* encodes the lysine secretion system LysE in *C. glutamicum* (Vrljic *et al*., 1996) and its deletion is a common strategy used to enhance L-lysine-derived compounds production (Pérez-García and Wendisch, 2018), including 5AVA (Rohles *et al*., 2016). However, in the present study, L-lysine intracellular availability was increased by heterologous expression of the *E. coli* permease gene *lysP* (Ruiz *et al*., 2011).

## 5. Conclusion

In conclusion, *avaC* from *C. intestinalis* enabled valerolactam production in *C. glutamicum* and proved to be a promising alternative to previously described lactam-forming enzymes. Process performance was strongly influenced by oxygen availability and carbon supply, with 50% rDO and carbon-limited fed-batch cultivation providing the most favorable balance between growth, productivity, yield, and by-product formation. These findings establish an AvaC-based route for valerolactam biosynthesis in *C. glutamicum* and provide a basis for further improvement through combined metabolic engineering and bioprocess optimization.

## Conflicts of Interest

The authors declare no conflict of interest.

## Author Contributions

**Henriette Victoria Bostad**: investigation, writing - review and editing. **Luciana Fernandes Brito**: investigation, writing - original draft, writing - review and editing, methodology, conceptualization. **Fernando Pérez-García**: investigation, writing - original draft, writing - review and editing, supervision, project administration, methodology, validation, conceptualization, funding acquisition.

## Funding

Fernando Peréz-García was funded by The Research Council of Norway within the FRIPRO funding scheme (project number 345245). Luciana Fernandes de Brito was funded by the Novo Nordisk Foundation (Grant number NNF24OC0094177).

## Supporting information

Supplementary material is provided

## Acknowledgements

Not applicable.

## Notes

### Competing Interest Statement

The authors have declared no competing interest.

