## Supplementary material for "A novel route for valerolactam production in *Corynebacterium glutamicum*: metabolic engineering and bioprocess optimization"

**Table S1.** Primers used in this study.

| Gene | Template | Primer Name | Primersequence5'→3' | Description |
|---|---|---|---|---|
| *davB* | *P. putida kt2440* gDNA | davB-fw1 | _AGACCATGGAATTCGAGCTCGGTACCCGGG_GAAAGGAGGCCCTTCAG**ATGAACAAGAAGAACCGCCAC** | Forward primer for amplification of *davB* |
| | | davB-rv | **TCAATCCGCCAGGGCGATCGG** | Reverse primer for amplification of *davB* |
| davA | *P. putida kt2440* gDNA | davA-fw | _GAAATCGGCCCGATCGCCCTGGCGGATTGA_GAAAGGAGGCCCTTCAG**ATGCGCATCGCTCTGTACCAG** | Forward primer for amplification of *davA* |
| | | davA-rv | _TGCATGCCTGCAGGTCGACTCTAGAG_**GATCTCAGCCTTTACGCAGGTGCAG** | Reverse primer for amplification of *davA* |
| *davB* | *P. putida kt2440* gDNA | davB-fw2 | _ATGAAGTTCCCGCAGAACGATAAGAAATAA_GAAAGGAGGCCCTTCAG**ATGAACAAGAAGAACCGCCAC** | Alternative forward primer for *davB* that creates a Gibson overlapping region with *lysP* |
| *lysP* | *E.coli MG1655* gDNA | lysP-fw | _AGACCATGGAATTCGAGCTCGGTACCCGGG_GAAAGGAGGCCCTTCAG**ATGGTTTCCGAAACTAAAACC** | Forward primer for amplification of *lysP* |
| | | lysP-Rv | **TTATTTCTTATCGTTCTGCGG** | Reverse primer for amplification of *lysP* |
| *avaC* | Codon optimized synthetic *avaC* | avaC-fw | _GAGCTGCACCTGCGTAAAGGCTGAGATCCT_GAAAGGAGGCCCTTCAG**ATGGGTAAAAAATATGCAATCG** | Forward primer for amplification of *avaC* |
| | | avaC-Rv | _CAAGCTTGCATGCCTGCAGGTCGACTCTAG_**TTATTCGGAGTACTTGTTGCC** | Reverse primer for amplification of *avaC* |

Bold letters: primer annealing sequences; underlined letters: ribosomal binding site sequences; italicized letters: Gibson assembly-directed overlapping sequences.

### Codon optimized *avaC* sequence:

ATGGGTAAAAAATATGCAATCGTTGGTGGTAAGTTGATCGATGGCACGGGTGCGGATCCAGT
TGAAAATTCTTTGGTACTTGTAGATGAAAACGGCAAGATCGAGTACGCGGGTGCACACCAAG
ACACGCCGGAGGGCTACGAGGTTATTGATGCGACCGGTAAAACGGTCATGCCGGGTCTGATC
GATACCCACCTTCATTTTAGCGGCAACTTGACCGATGATGATACTGACTGGGTGATGCAGCC
GTTGCTGGAGAAGCAAGCCGTCGCAGTCAAACAGGCATACGATTGTCTGACCCACGGCTTGA
CGACGGTGTGCGAAATCGGTCGTTTCGGCATCCAGATTCGTGATTGTATTGACAAAGGTGTT
TTCAAAGGTCCCCGTGTTCTCGCTACTGGATTGGGCTTCTGCCGCACCGCGGGACACGGTGA
TTCCCACCACTGTTCCCAGTTGGAAAACAAGGAGTCCCACCCTTGGGGAGACCAAGTCGATG
GTCCTTGGGATCTGCGTAAAGCTGTTCGCCGCCGGCTCCGTGAGAACCCAGACGCTATCAAA
ATCTGGGCAACCGGCGGCGGCATCTGGCGCTGGGACTCAGGCCGCGATCAGCATTACTGTTC
GGAAGAGATTCAGGCCGTTATTGACGAGGCTAAACTGGTAGGTATCCCTGTTTGGTCGCACT
GCTACAACAACCATGCGGCTGCGTATGATTCAGTCCGTTTTGGATGCGAACAACTCATCCAT

GGTTTCGACATTGACGAACGTACTATGGATCTCATGGCCGAACAGGGCACCTTTTTCACGCC
CACTATCGCATTCCTGCCGACTTGGTACTCCACTTACCCGCCGGTCTATGTGCCGGAGCTGC
ACGATAAGTACGAGGGCACTCTGGTCGAAAAGGAGCTTCAACGCAACTACGATTGTCTCCGT
GAGGCAAAAAAGCGCGGTGTTGTGATGACCATCGGATCAGATTCCTTTAGCTTTGTGACCCC
GTATGGTACTTGCTCGATCGAAGAAATGTACGAGTTCGTTGATAAGATCGGCTTCACCCCCG
TGGAAACCATTACTTGCGCCACCCTGAACGGCGCGAAAATGTGCCACATCGAGGATGAGACC
GGATCCCTGGAAGCAGGCAAGTGCGCCGACCTTCTCGTGGTCAACGGCGATGTGGCAGCTGA
CATCCACACCCTTAACGTCGATAACATGGACGTGATCATGAAGGATGGATGGATCGTGGACG
CTGGTACCTTCGGAGAAGGCAACAAGTACTCCGAATAA
